# Multiomic profiling of hypoxic glioblastoma stem cells reveals expansion of subpopulations with distinct epigenetic and CNV profiles

**DOI:** 10.1101/2025.05.05.652238

**Authors:** Adrianne Corseri, Travis Moore, Nicole Szczepanski, Hyeyeon Hwang, Alan Zdon, Gurkan Yardimci Galip, Nikos Tapinos

## Abstract

Glioblastoma is characterized by extensive intratumoral heterogeneity driven by dynamic microenvironmental cues such as hypoxia. While transcriptional and epigenetic variability have been separately linked to hypoxia responses, the integrated impact of hypoxia on gene regulation and clonal architecture in glioblastoma stem cells (GSCs) remains poorly defined. We applied singlenucleus multi-omics—integrating RNA-seq and ATAC-seq—to patient-derived GSCs cultured under normoxic or hypoxic conditions. This enabled simultaneous profiling of gene expression and chromatin accessibility within the same cells. Transcription factor (TF) regulatory networks were inferred using Dictys, while RNA-chromatin dynamics were modeled with MultiVelo. Clonal structure and copy number variations (CNVs) were resolved at single-cell resolution using RIDDLER on snATAC-seq data. Hypoxia induced the emergence of four distinct GSC subpopulations with unique transcriptomic and epigenetic profiles enriched for mesenchymal, angiogenic, and proliferative signatures. Regulatory network modeling revealed novel hypoxia-associated TFs—SP2, CREM, and ETV3—that modulate downstream oncogenic pathways. Trajectory analysis uncovered hypoxia-driven reversals in RNA-chromatin coupling, revealing dysregulated future transcriptional states of key genes such as MMP16 and SVIL. CNV profiling identified 13 clonal substructures, with specific clones (e.g., 5, 6, 9) selectively enriched under hypoxia and harboring distinct chromosomal alterations. These results demonstrate coordinated remodeling of GSC gene regulation and clonal fitness in response to hypoxic stress. Our findings reveal that hypoxia drives concurrent epigenetic, transcriptomic, and clonal selection in glioblastoma stem cells. This integrated model of hypoxia-induced plasticity provides mechanistic insights into tumor adaptation and identifies novel regulators that may serve as targets for therapeutic intervention in the hypoxic niche of glioblastoma.

## Introduction

Glioblastoma is the most common, aggressive and lethal form of primary brain tumors in adults marked by profound inter- and intra-tumoral heterogeneity, treatment resistance, and inevitable recurrence.^1, 2^ Standard treatment for glioblastoma consists of surgical resection, radiation, and adjuvant temozolomide therapy.1 Despite incremental advances in treatment, median survival after diagnosis is under 2 years.2 Among the key drivers of heterogeneity in glioblastoma is the hypoxic tumor microenvironment, a hallmark of high-grade gliomas.^3^ Hypoxia not only limits therapeutic efficacy but also promotes stem-like phenotypes, immune evasion, and cellular plasticity.^4–6^ Despite its recognized role in shaping transcriptional states and influencing cell fate decisions, the extent to which hypoxia exerts selective pressure to rewire the epigenetic landscape and promote the expansion of distinct cellular subpopulations remains incompletely understood.

Glioblastomas are also characterized by a complex genomic architecture dominated by copy number variations (CNVs), and structural rearrangements, which drive tumor evolution and therapy resistance.^7, 8^ Although CNVs and structural variants are well-documented contributors to glioblastoma progression, their interplay with dynamic environmental cues such as hypoxia and the resultant impact on epigenetic remodeling has not been systematically explored. Recent advances in spatial and single-cell multiomic technologies have enabled high-resolution mapping of tumor ecosystems, revealing spatially segregated transcriptional programs and lineage hierarchies.^9–11^ However, a comprehensive interrogation of how hypoxia shapes the transcriptomic, epigenetic, and CNV landscape at single-cell resolution remains lacking.

Our previous work has demonstrated that microenvironment pressure and chromatin remodeling, particularly through the influence of ChI3L1^12^ and epigenetic modulators such as HDAC7^13^ respectively, play a critical role in orchestrating transcriptional programs and phenotypic plasticity in glioblastoma. Building upon these findings, we now investigate how hypoxia drives the emergence of phenotypically distinct subpopulations through coordinated transcriptomic, epigenetic, and structural alterations.

Here, we perform integrated single-cell multiomic profiling of hypoxic versus normoxic patient-derived glioblastoma stem cells to define how oxygen deprivation acts as a selective force to remodel chromatin accessibility and expand subclonal populations with distinct CNV and epigenetic profiles. Our findings uncover a previously unappreciated role for hypoxia in exerting evolutionary adaptive pressure to cancer stem cells and highlight a potential convergence of environmental and genetic factors in shaping glioblastoma heterogeneity.

## Results

### Identification of hypoxic subpopulations of cells in GSCs using multi-omics

To elucidate the epigenetic mechanisms underpinning the aggressive phenotype of glioblastoma (GBM) under hypoxic conditions, we performed integrative transcriptomic and chromatin accessibility analyses. Primary GSCs were cultured under normoxic or hypoxic conditions, then processed and sequenced using single-nucleus multi-omics. This approach enabled simultaneous capture of gene expression (from mRNA) and chromatin accessibility (from ATAC) within the same cells, for both conditions (Fig. 1a).

**Fig. 1.**
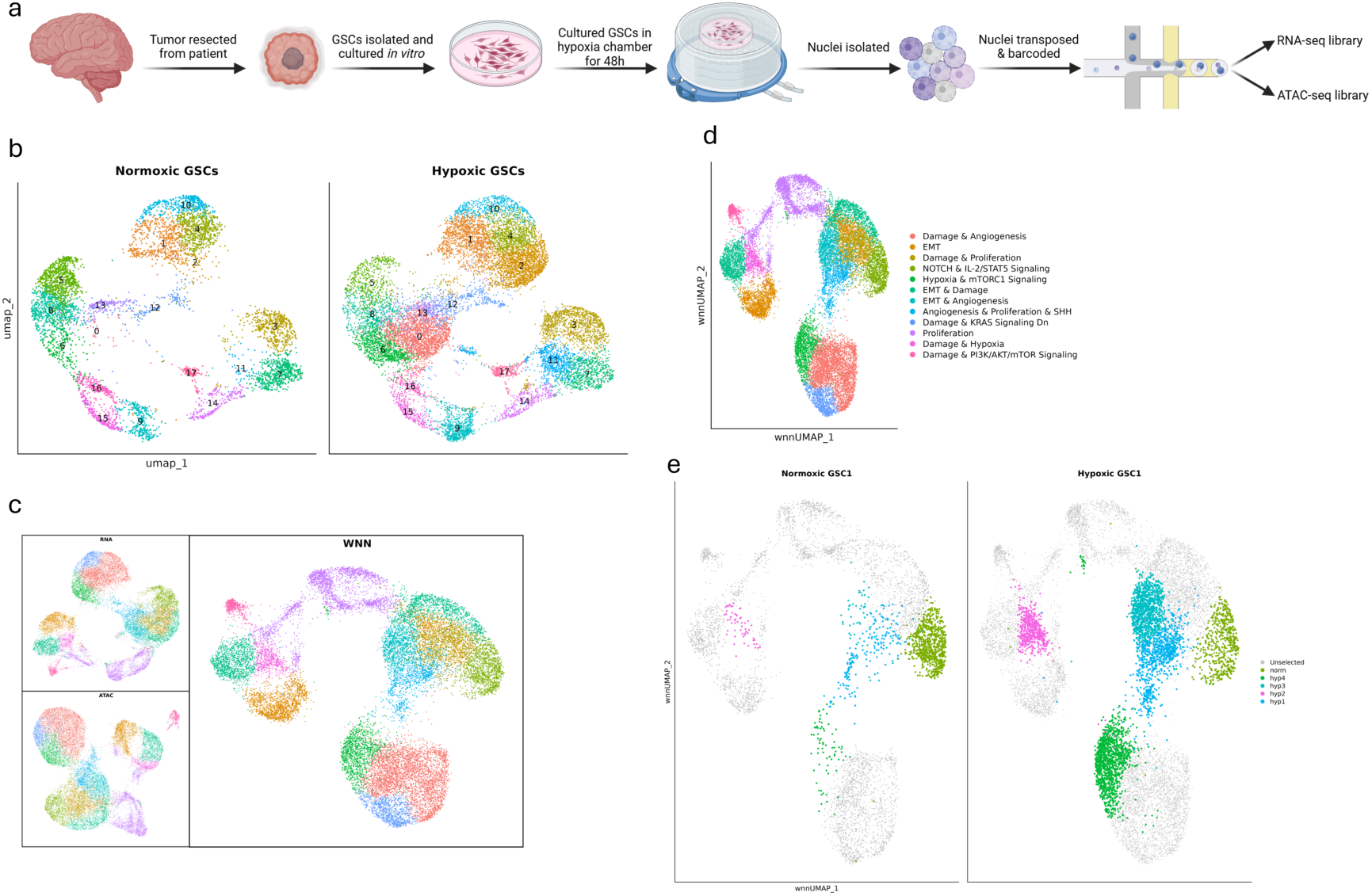
Identification of hypoxic subpopulations of cells using multi-omics. **a.** Schematic of GSC isolation, *in vitro* culture, and multi-omic sequencing. **b.** UMAP showing the differences between normoxic and hypoxic GSCs, in sn-RNA-seq only. **c.** UMAP showing optimized clustering across all cell types, using gene expression (RNA), chromatin accessibility (ATAC), or both (WNN). **d.** Cell signature markers for each cluster. **e.** Highlighting the different unique cell populations between conditions using the integrated multi-modal data. **f.** Multiome velocity, showing future states of the cells.

Next, we employed the Jaccard similarity index^14^ to evaluate and optimize clustering parameters, ensuring stable cell clusters. The resulting clustering allowed us to identify three GSC subpopulations that are present in the hypoxic cell population, but largely absent from the normoxic sample (Fig. 1b). To further refine clustering, we integrated ATAC-seq and transcriptomic profiles via a weighted nearest neighbor UMAP framework,^15^ resulting in a more comprehensive representation of cellular states (Fig. 1c). Then, we assigned functional cell signature markers to each cluster based on enrichment of hallmark gene sets from the MSigDB Hallmark collection^16^ (Fig. 1d). This integrative approach revealed four hypoxia-specific subpopulations and one distinct normoxia-specific subpopulation (Fig. 1e), an expansion upon the three previously found from analysing just the RNA-seq without the chromatin accessibility data. These distinct clusters form the basis for subsequent analyses.

### Characterization of Condition-Specific GSC Subpopulations

To better understand the biological relevance of the hypoxia- and normoxia-specific clusters, we first analyzed the transcriptomic profiles of all the clusters in the fully integrated sample (Fig. 2a). Hypoxia-specific clusters were predominantly associated with mesenchymal-like transcriptional programs, consistent with the classification framework from Neftel et al.^17^ (Supp. Fig. 2). Conversely, the normoxic cluster exhibited signatures of multiple neural lineages, including NPC-like (neural progenitor cells), OPC-like (oligodendrocyte progenitor cells), and AC-like (astrocyte) states. The ‘Proliferation’ cluster (a combination of clusters 11, 12, 14, 15) is highly aligned with the cell cycling markers previously identified. Remaining clusters, shared across both oxygen conditions, displayed mixed subtype markers.

**Fig. 2.**
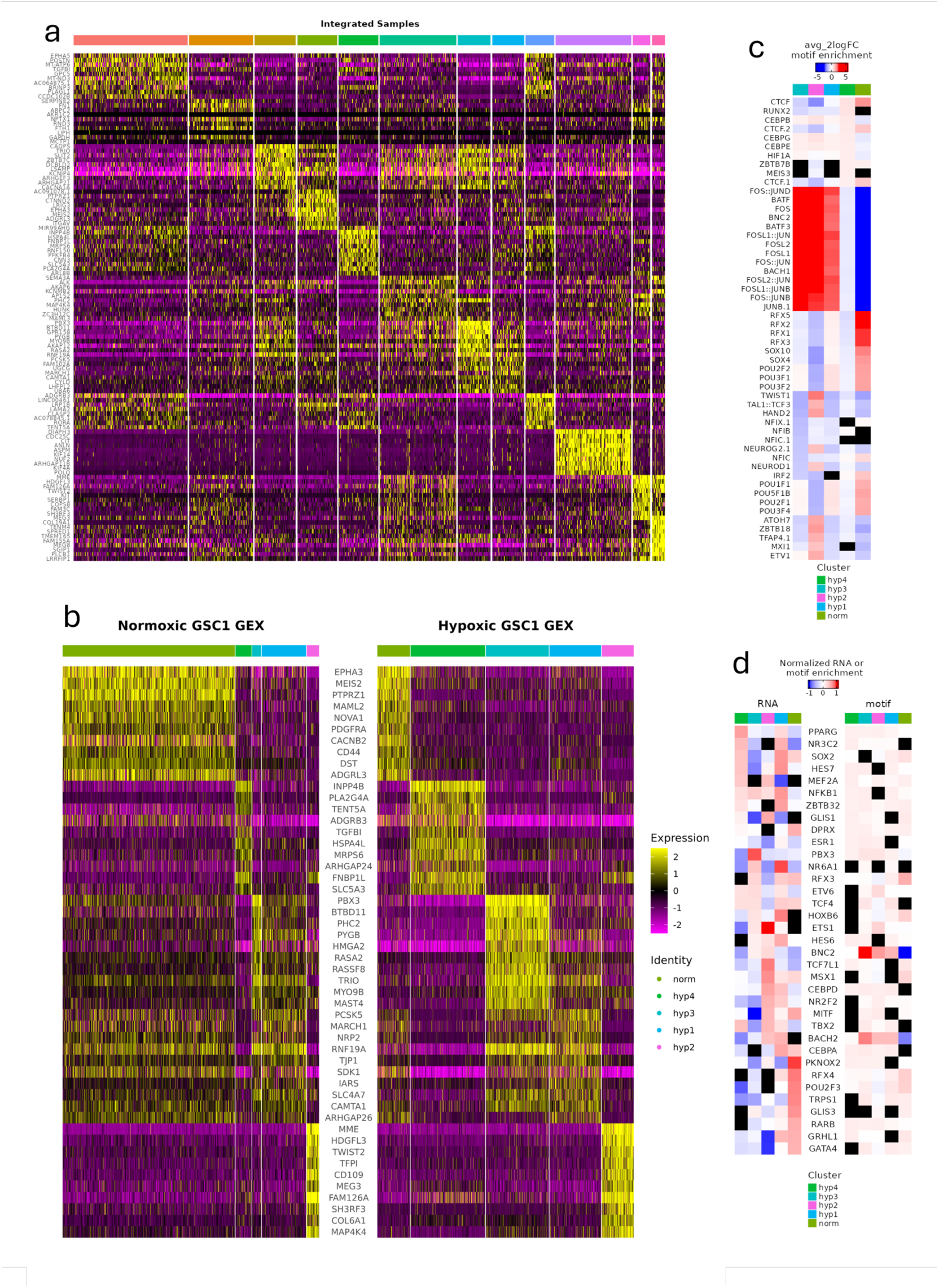
Preliminary analysis of transcriptome and epigenome of each condition. **a.** Heatmap of top 10 markers of each cluster in the integrated dataset. **b.** Top 10 gene markers the unique clusters, compared between conditions. **c.** Top motifs, by motif enrichment score, in each unique cluster of interest. **d.** Normalized gene expression & motif enrichment for top 10 active Transcription Factors in the unique clusters.

Given the heterogeneity of the dataset, we focused on the 4 clusters with clear hypoxia-specific enrichment: Hypoxia & mTORC1, EMT & Angiogenesis, Damage & Hypoxia, Angiogenesis & Proliferation & SHH, as well as the cluster specific to normoxia, NOTCH & IL2-STAT5. We first analyzed the top differentially expressed genes within these clusters (Fig. 2b, Supplementary Table 5). All of these have already been identified in glioblastoma, (e.g. TRIO,^18^ PBX3^19^), in hypoxia (e.g. AKAP12^20^), or in both (INPP4B,^21^ TWIST2,^22, 23^ MEIS2.^24, 25^ Given that none of the top genes in the transcriptome are novel to GBM or hypoxia, we aimed to extract insights uniquely enabled by the multi-omic approach.

### Identification of transcription factor regulatory networks in hypoxic glioblastoma stem cells

As transcription factors (TFs) are central to gene regulation, we next investigated TF motif enrichment in the chromatin profiles of each cluster (Fig. 2c). While this revealed potentially relevant motifs, the overlap between enriched motifs and transcriptionally active genes was limited (Fig. 2d), suggesting that further modeling was necessary to capture regulatory dynamics. To identify candidate regulatory TFs, we used Dictys,^26^ an approach that builds a TF binding network and further refines it using transcriptomic data. This enabled us to extract TF activity specific to each cluster for both conditions. Several TFs showed strong regulatory influence within hypoxia-specific clusters (Fig. 3a), including some not previously identified in glioblastoma. The first was SP2, part of the SP/KLF family with conserved zinc finger domains,^27^ thought to be essential in certain biological processes, including cell cycle genes and VEGF.^28^ The second is CREM, the cAMP responsive element modulator, which has been linked to reducing growth and invasion in glioblastoma.^29^ The third is ETV3, part of the ETS family of TFs, known to be activation targets of the Ras-MAP kinase pathway,^30^ an important signaling pathway involved in glioma proliferation, angiogenesis, and survival.^31^ Furthermore, cluster-specific regulation is unveiled using this tool, allowing us to look at both the top activated and top inactivated downstream gene targets of these TFs (Fig. 3b). A number of TFs that are already well-established in glioblastoma and associated cancers (e.g. SALL4,^32, 33^ MYC,^34^ FOS,^35, 36^ JUNB^37, 38^) are revealed to be targets of the identified targets. It also highlights non-TF gene targets such as NR2C1 and NFIC, which may represent unexplored drivers of GSC phenotype under hypoxic stress.

**Fig. 3.**
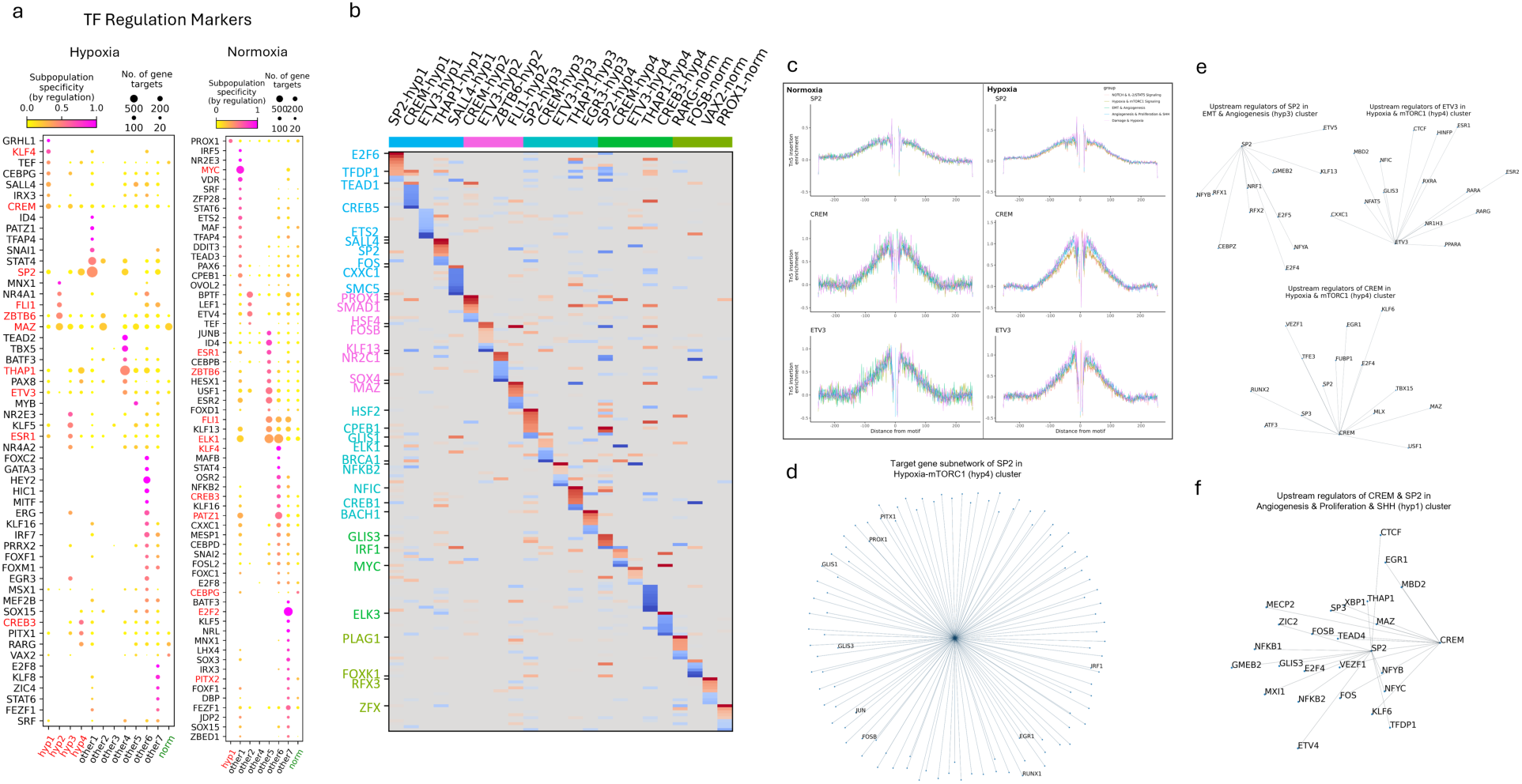
Transcription Factor regulatory networks in hypoxic GSCs. **a.** Transcription factor regulation marker discovery across both conditions, based on target count and and specificity. **b.** Regulation heatmap in hypoxic sample from each TF marker in its corresponding cluster, of top ten most or least expressed targets. **c.** TF footprinting of SP2, CREM, and ETV3. **d.** Target gene subnetworks of SP2, in the Hypoxia & mTORC1 Signaling (“hyp4”) cluster. **e.** Upstream regulators of SP2, CREM, and ETV3 in select clusters of interest. **f.** Upstream regulators of both SP2 and CREM in the Angiogenesis & Proliferation & SHH (“hyp1”) cluster, indicating the link between the two.

After identifying a few transcription factors that had not been previously linked to glioblastoma, we wanted to confirm they play a role in hypoxic GSCs, as well as further elucidate their characteristics and the role they play. Motif footprinting of SP2, CREM, and ETV3 (Fig. 3c) displays enrichment of Tn5 integration surrounding their corresponding motif sites, with particularly stable footprints in the hypoxic samples, further supporting the data found in *Dictys*. We also observed that average expression is higher in hypoxia than in normoxia, and increasing over time (Supp. Fig. 5), except for CREM, which is lower in hypoxia. We built a target gene subnetwork (Fig 4d) for these transcription factors of interest, expanding upon the previous target gene heatmap. We see that a number of targets within all these subnetworks include transcription factors and genes already well associated with glioblastoma, and upstream regulators of these identified TFs (Fig. 3e) also exhibit key drivers of glioblastoma, further supporting the idea that these are important in driving the mesenchymal phenotype. Additionally, SP2 is one of the identified upstream regulators of CREM (Fig. 3f), indicating a possible signaling pathway, including the possibility of a metabolic pathway operating during hypoxia, as CREM is one of the main regulators of PKA (alongside CREB), and PKA phosphorylates many metabolic enzymes.^39^

**Fig. 4.**
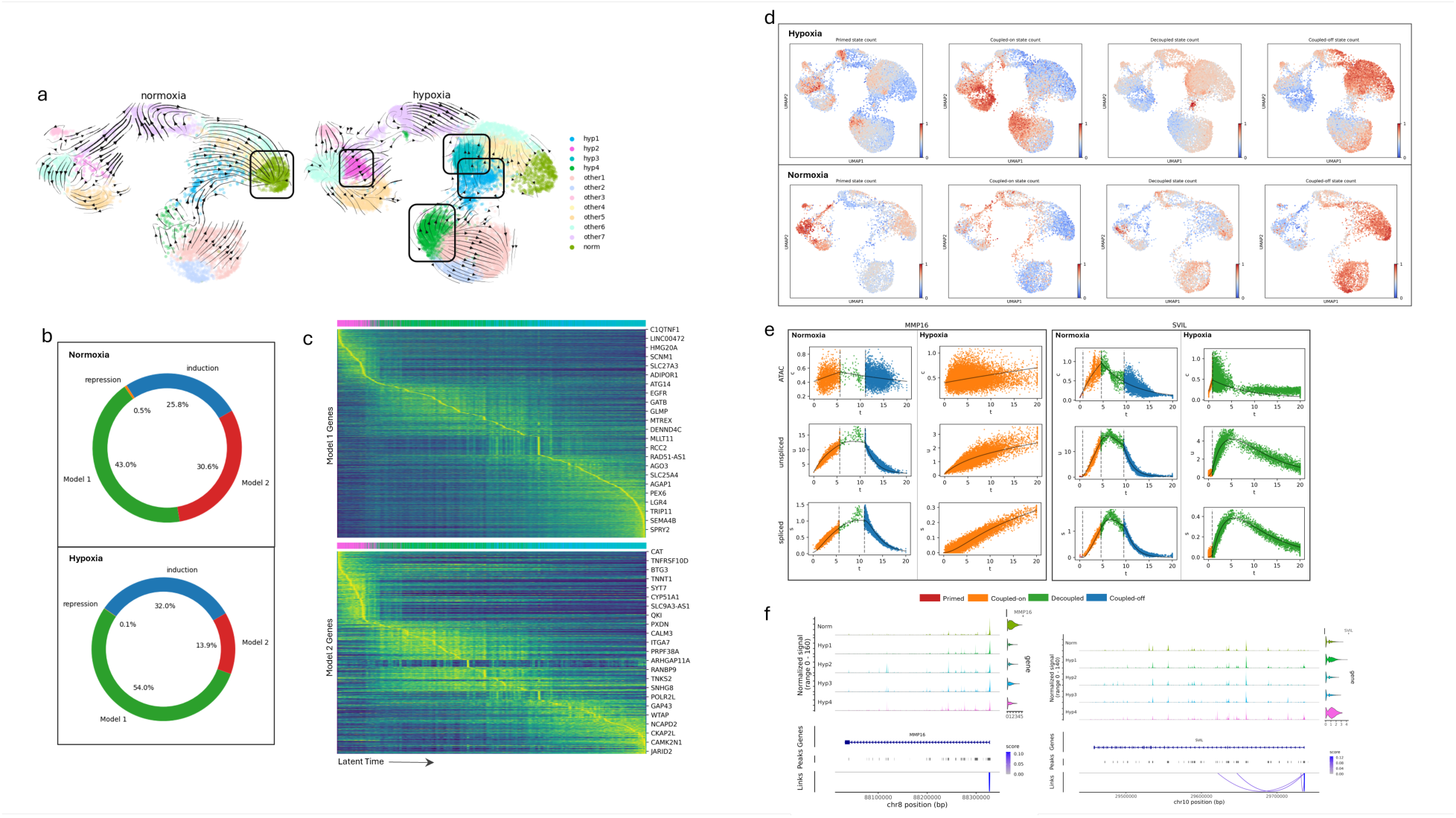
Hypoxic-induced transcriptomic and epigenetic cellular states. **a.** Multiome velocity of both conditions. **b.** Proportion of all included genes across each possible fit model (Model 1, Model 2, induction-only, repression-only) in both normoxia and hypoxia. **c.** Smoothed spliced counts heatmaps of model 1 and model 2 genes in just the hypoxia-specific clusters, as a function of latent time. **d.** UMAP assigning each cell to one of the four fit states, across both conditions. **e.** Changes in gene dynamics between the two conditions for two potential gene targets, MMP16 and SVIL. **f.** Linked peaks to genes, for MMP16 and SVIL.

### Characterization of hypoxia induced transcriptomic and epigenetic cellular states

To gain deeper insight into the temporal regulatory processes shaping condition-specific phenotypes, we employed MultiVelo^40^ to predict cell state trajectories. MultiVelo is a framework that models RNA velocity in conjunction with chromatin accessibility, using a probabilistic latent variable model to estimate the switch time and rate parameters of chromatin accessibility and gene expression. We observe pronounced epigenomic and transcriptomic velocity field changes following exposure of GSC to hypoxia (Fig. 4a), suggesting hypoxia-induced cell state transition changes. There are striking directionality changes in the multiome velocities in the “Angiogenesis & Proliferation & SHH” cluster (hyp1), “Damage & Hypoxia” cluster (hyp2), “EMT & Angiogenesis” cluster (hyp3) and “Hypoxia & mTORC1” cluster (hyp4). In all four clusters we notice complete reversal of the cell state trajectories suggesting a prominent role of the hypoxic microenvironment in shaping epigenomic and transcriptomic cellular states of GSCs, further confirming these clusters as essential in driving hypoxia in GSCs.

Beyond predicting future transcriptional states, MultiVelo enabled us to categorize genes into mechanistic models of activation and repression (Fig. 4b). Specifically, it distinguishes between two “on” gene models: Model 1, where chromatin closure precedes transcriptional repression, and Model 2, in which transcriptional repression occurs prior to chromatin closure. Using this framework, we identified Model 1 and Model 2 genes for each condition (Supp. Fig. 3a and 3b). There were further refined to represent these genes within each unique cluster under hypoxic (Fig. 4c) and normoxic (Supp. Fig. 3e) conditions, revealing distinct regulatory regimes (Supp. Table 1 and 2). MultiVelo also allowed us to identify “fit states” for each gene, over time: primed, coupledon, decoupled, and coupled-off (Fig. 4e). These states indicate the relationship between chromatin accessibility and gene expression: one is on while the other is off, both are on and rising, one of the two is very low/turned off, and both are off, respectively. This allowed us to quantify the number of genes in each transcriptional state per cell across both environments, highlighting broad regulatory shifts that may underlie hypoxia-induced phenotypic changes.

Multivelo’s fit states allowed us to identify any genes that appeared to have highly changed expression patterns during hypoxia. Any genes that appeared to have opposite effects, or a reversed fit state (Supp. Table 3 and 4), were considered to be additional potential targets. Among these, we found some that have already been shown to have effects in GBM or hypoxia, such as AKAP12,^20, 41^ CREB5,^42, 43^ and TWIST2.^22^ Of those that exhibited unexpected future behavior and were not already known to GBM or hypoxia, most showed just small amounts of gene expression changes, and were deemed uninteresting. In the end, two stood out: MMP16 and SVIL. MMP16 is a one of many matrix remodelling proteins, and has been linked invasion and proliferation.^44, 45^ SVIL has been linked to angiogenesis in liver cancer,^46^ suppression of p53, enhancing cell survival,^47^ and even progressing ovarian cancer and epithelial-mesenchymal transition.^48^ These factors all indicate promising targets for future glioblastoma treatments. We see that despite lower average expression over time (Fig. 4e) in the hypoxic cells, there is continuously increasing expression long past the ending point for the same gene in the normoxic sample, indicating dis-regulation, as they are never turned off. We also find strong links between these genes and the associated peaks in the surrounding chromatin region (Fig. 4f), providing possible regulatory elements (most likely promoters given the localization, but possibly enhancers) for these genes.

### Clonal heterogeneity of glioblastoma stem cells is altered under hypoxic conditions

Glioblastoma cells are known to exhibit high levels of copy number variation (CNV), which can potentially drive gene dosage effects altering TF expression and other chromatin remodeler protein complexes, leading to gross gene mis-regulation, transition to mesenchymal subtype and other alterations, potentially under hypoxic conditions. To study such potential CNV driven effects and CNV selection between different conditions, we identified single-cell resolution CNV events using the snATAC-seq normal and hypoxic samples using the RIDDLER algorithm.

RIDDLER identified a rich clonal heterogeneity structure with thirteen distinct CNV clusters identified via an unsupervised clustering algorithm. In Figure 5a, we visualized the genome wide CNV profile of each single cell in a heatmap after clustering and overlaying each single cell with associated functional pathway labels and normoxic or hypoxic conditions. While specific CNV clusters are enriched for pathway annotations, each clone is visually represented by both hypoxic and normoxic cells, suggesting a lack of dramatic selection of clonal signatures under the hypoxic conditions. However, due to the short treatment of glioblastoma stem cells under hypoxia, we performed statistical hypothesis testing asking whether any clonal cluster is underrepresented or overrepresented in each condition using a chi-square test. These tests revealed that clonal clusters 1 and 3 are significantly enriched in the normoxic cells, whereas clones 5, 6, 9 are depleted in the normoxic conditions (Fig 5b). Taken together, these results indicate that glioblastoma stem cells undergo clonal selection when placed in a hypoxic environment. We further investigated whether specific regions of the genome exhibit differential CNV events. By performing a Wilcoxon rank sum test over the CNV state of each genomic window, we identified specific regions of each clone that have a statistically significant differential CNV state. In Fig 5c, heatmap visualization of specific regions on chromosomes 10, 14, and 15 are shown with CNV states distributions across the two conditions. Our results demonstrate that the rich CNV diversity across glioblastoma stem cells are affected by hypoxic microenvironment, and we can pinpoint such regions through computational identification of CNV states at megabase resolution.

**Fig. 5.**
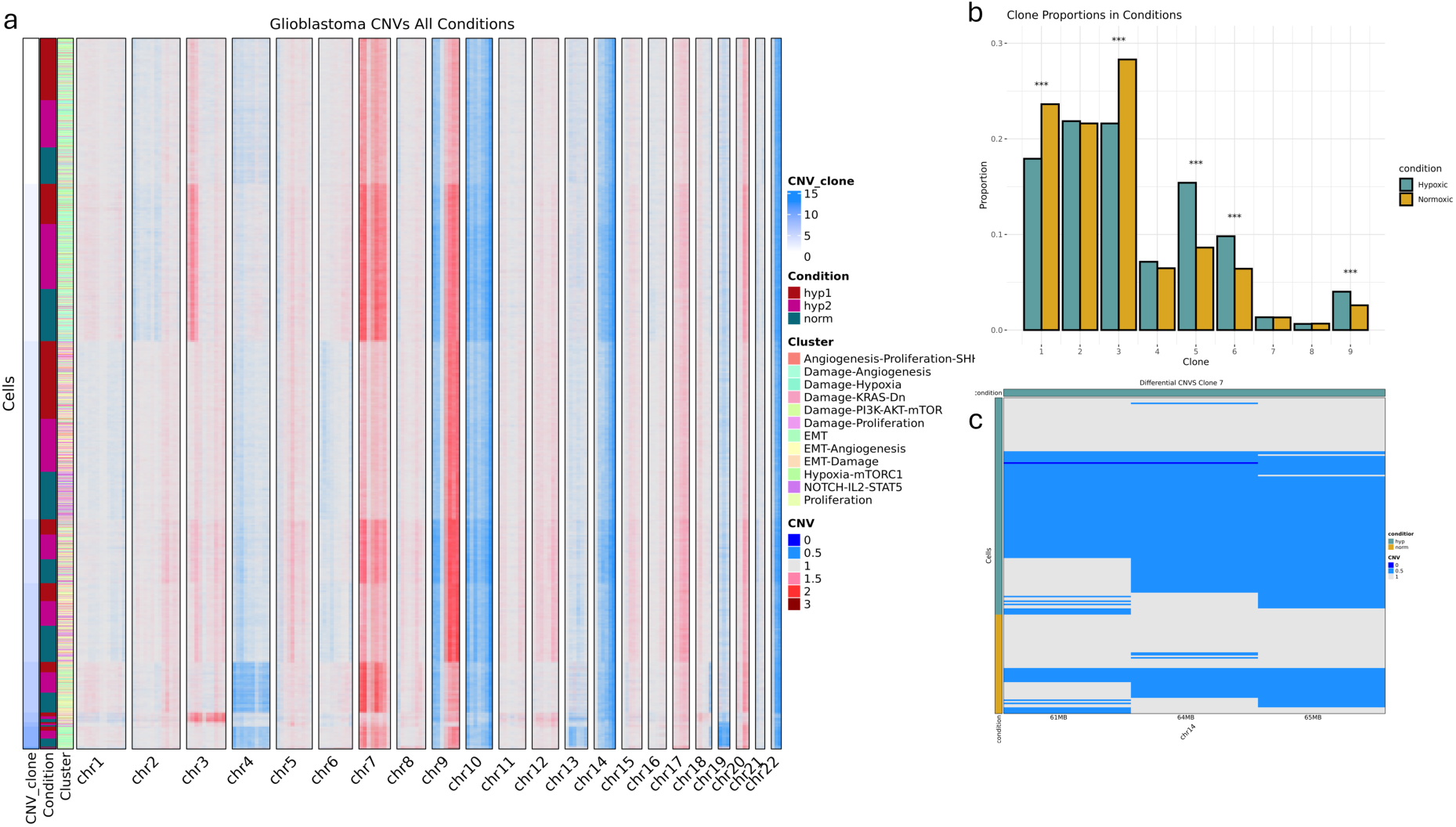
Clonal heterogeneity of glioblastoma stem cells is altered under hypoxic conditions. **a.** Single-cell CNVs detected by RIDDLER for all cells in all conditions at 1MB resolution, ordered by CNV subclone (rows). Clones were defined by clusters found by Louvain clustering of CNVs. Copy number for each 1 MB window of each cell is colored by blue tones for losses, red tones for gains, and grey for neutral. Experiment condition and cell cluster membership are also included on the row annotations. 13 total clonal group were identified. **b.** Proportion of cells in hypoxic and normoxic conditions for major subclones. indicates significant difference (p<0.001) from chi-squared test of clone proportion between conditions. **c.** Heatmaps of differential CNV windows between hypoxic and normoxic groups in clones with significant proportion differences. Differential windows were identified using a Wilcoxon Rank-Sum test with FDR corrected p-value of 0.05 or less and at least a 10% difference in CNV frequency. Column annotations indicate which condition group has a higher prevalence of CNVs for each window.

## Discussion

In this study, we leveraged integrative single-nucleus multiomic profiling to uncover how hypoxia remodels the transcriptomic and epigenetic landscape of glioblastoma stem cells (GSCs). By simultaneously capturing chromatin accessibility and gene expression within the same cells, we delineated four hypoxia-specific subpopulations characterized by distinct transcriptional programs, chromatin states, and trajectory dynamics (Figure 2a–d, Figure 3a–f). These findings extend prior work that described transcriptional heterogeneity in GSCs^17^ by demonstrating that hypoxia acts as a profound epigenetic pressure, driving the emergence of new cellular states with mesenchymal, angiogenic, and proliferative features.

Importantly, the identification of previously uncharacterized transcription factors (SP2, CREM, and ETV3) as key regulators within hypoxia-induced GSC subpopulations (Figure 3a–f) highlights novel regulatory networks that may underpin the aggressive behavior of hypoxic tumors. Our data suggest that these TFs orchestrate transcriptional programs that converge on established oncogenic drivers, such as MYC and FOS,^49, 50^ while also regulating genes such as NR2C1 and NFIC, whose roles in glioblastoma remain poorly defined. These observations nominate SP2, CREM, and ETV3 as potential therapeutic vulnerabilities within the hypoxic niche of glioblastoma.

The use of MultiVelo modeling revealed that hypoxia not only alters the steady-state expression of key genes but also rewires their future regulatory trajectories (Figure 4a–f). Specifically, we observed widespread reversals in cell state transitions and dynamic chromatin-transcriptional coupling in hypoxia-enriched clusters (Figure 4a). The identification of genes such as MMP16^51^ and SVIL,^46^ which exhibited prolonged dysregulated expression trajectories in hypoxia despite transcriptional downregulation (Figure 4e–f), underscores the complexity of hypoxia-mediated reprogramming and suggests that conventional steady-state analyses may underestimate the full extent of hypoxic remodeling.

To further understand the role of genomic instability in shaping these hypoxia-induced phenotypes, we performed single-cell CNV profiling using snATAC-seq data analyzed with RIDDLER. This revealed substantial clonal heterogeneity with 13 distinct CNV clusters. Although most clones were represented in both hypoxic and normoxic samples, statistical analysis showed that specific clones—particularly clones 5, 6, and 9—were enriched under hypoxia, while others (e.g., clones 1 and 3) were depleted (Figure 5b). These results suggest that hypoxia drives clonal selection, favoring subpopulations with distinct CNV profiles. Moreover, chromosomal regions such as 10, 14, and 15 exhibited condition-specific CNV alterations (Figure 5c), indicating that the hypoxic microenvironment not only remodels gene regulation but may also influence genomic stability and clone fitness.

Together, these data present a multi-layered model in which hypoxia exerts selective pressure across transcriptomic, epigenetic, and clonal dimensions, reshaping the functional landscape of GSCs. While previous work has explored some of these axes independently, our integrative approach provides a unified view of how the hypoxic niche promotes plasticity and aggressiveness in glioblastoma.

Several limitations warrant consideration. While our multiomic approach enables unprecedented resolution of hypoxia-induced states, functional validation of candidate TFs and regulatory elements is needed to confirm their causal role in GSC plasticity and tumorigenicity. Additionally, our study was conducted under controlled hypoxia in vitro; future efforts should aim to map these findings in patient-derived tumor samples in situ, incorporating the influence of additional microenvironmental factors such as immune cells and extracellular matrix components.^52^

In conclusion, our work reveals that hypoxia acts as a powerful driver of epigenetic and transcriptional diversification in glioblastoma, identifying novel regulators and providing a foundation for therapeutic strategies aimed at targeting the hypoxic niche. These findings highlight the importance of integrating multiomic and dynamic modeling approaches to fully capture the complexity of tumor evolution under microenvironmental pressures.

## Figures

**Supplementary Fig. 1.**
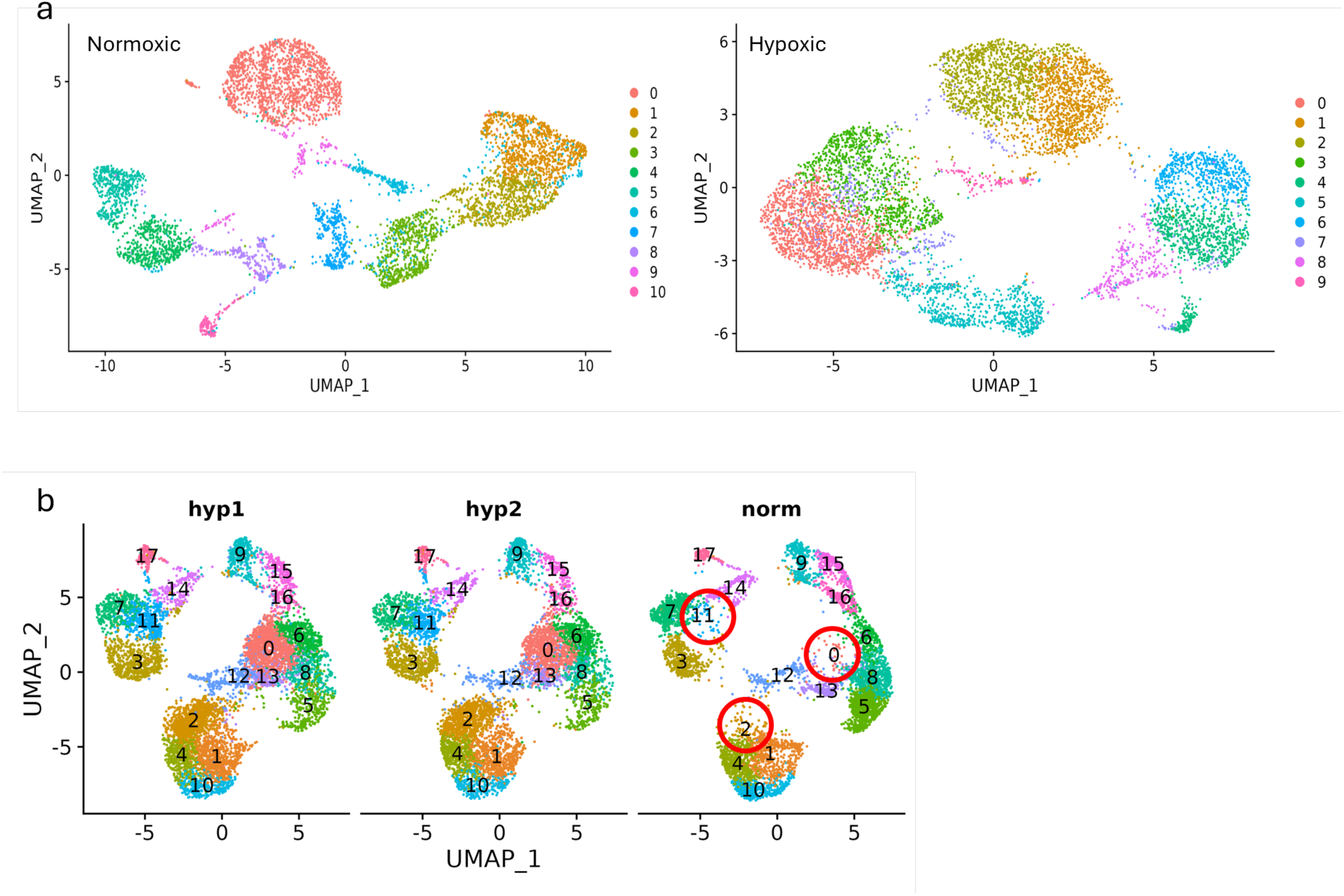
**a.** Default clustering parameters without integration on multiome samples. **b.** Jaccard optimized clustering from just RNA-seq data.

**Supplementary Fig. 2.**
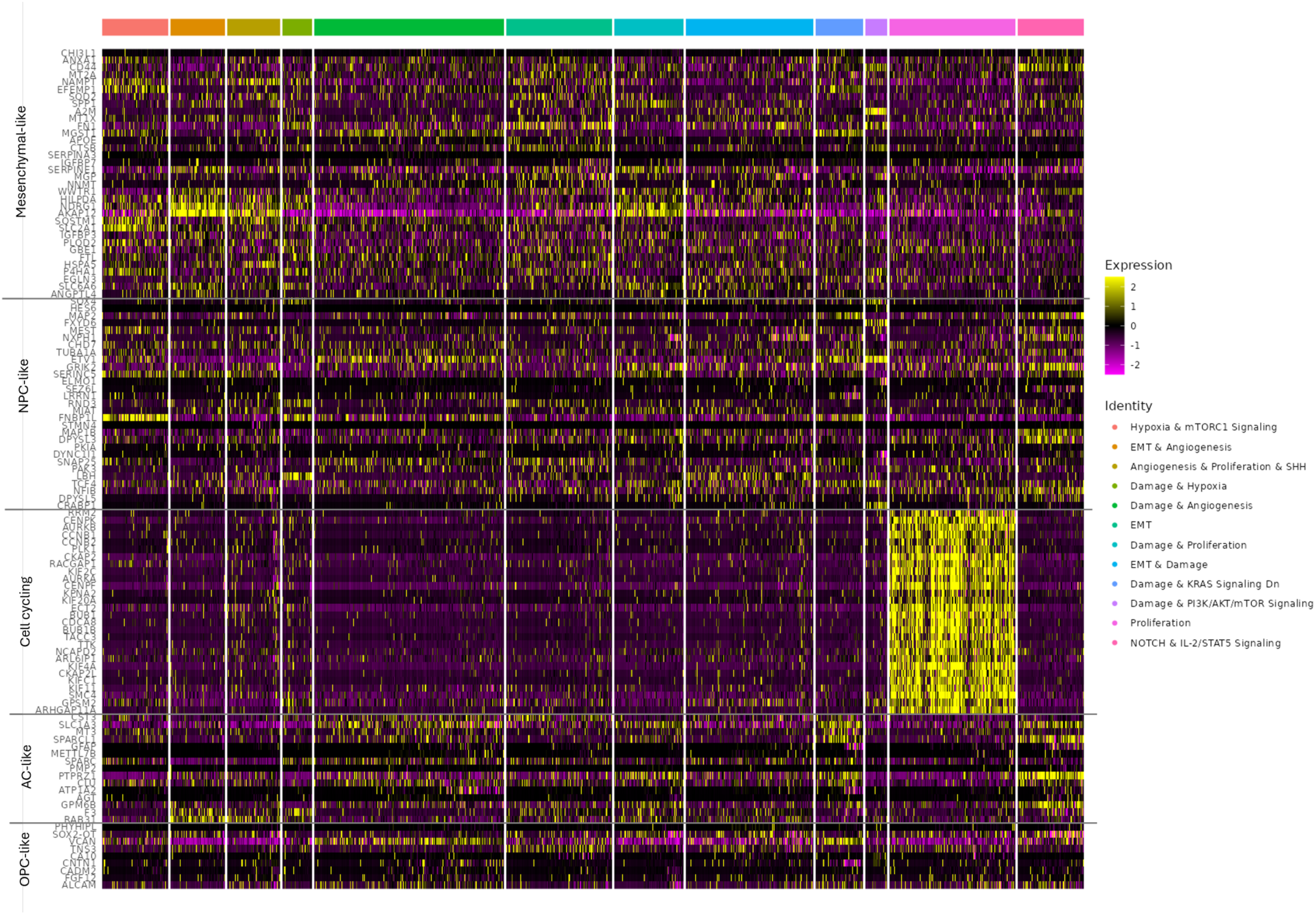
Classification of each cluster by molecular cell state, according to the Neftel^18^ model.

**Supplementary Fig. 3.**
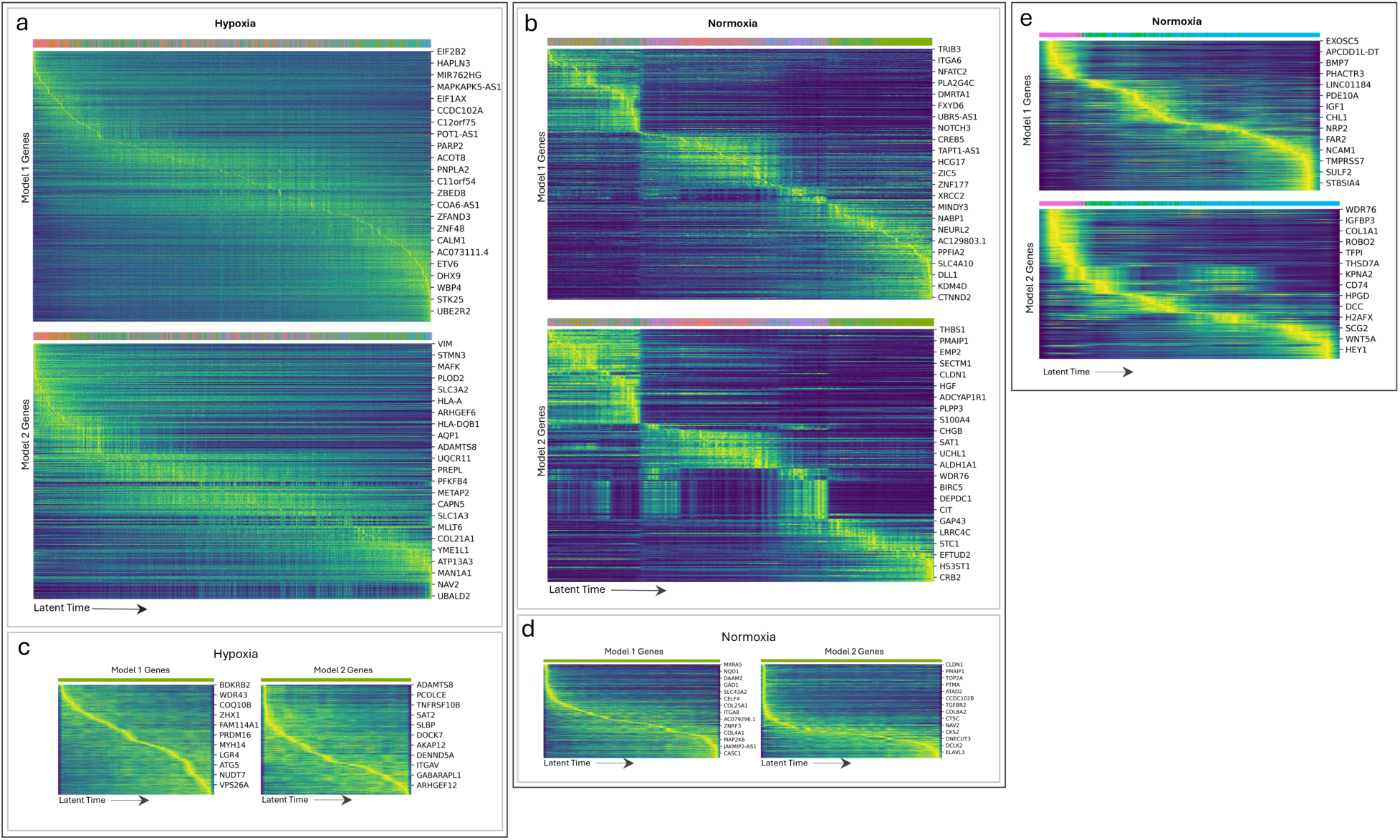
**a.** Model 1 and 2 genes across all clusters in the hypoxic sample. **b.** Model 1 and 2 genes across all clusters in the normoxic sample. **c.** Normoxic cluster cells in the hypoxic sample. **d.** Normoxic cluster cells in normoxic sample. **e.** Hypoxic cluster cells in the normoxic sample.

**Supplementary Fig. 4.**
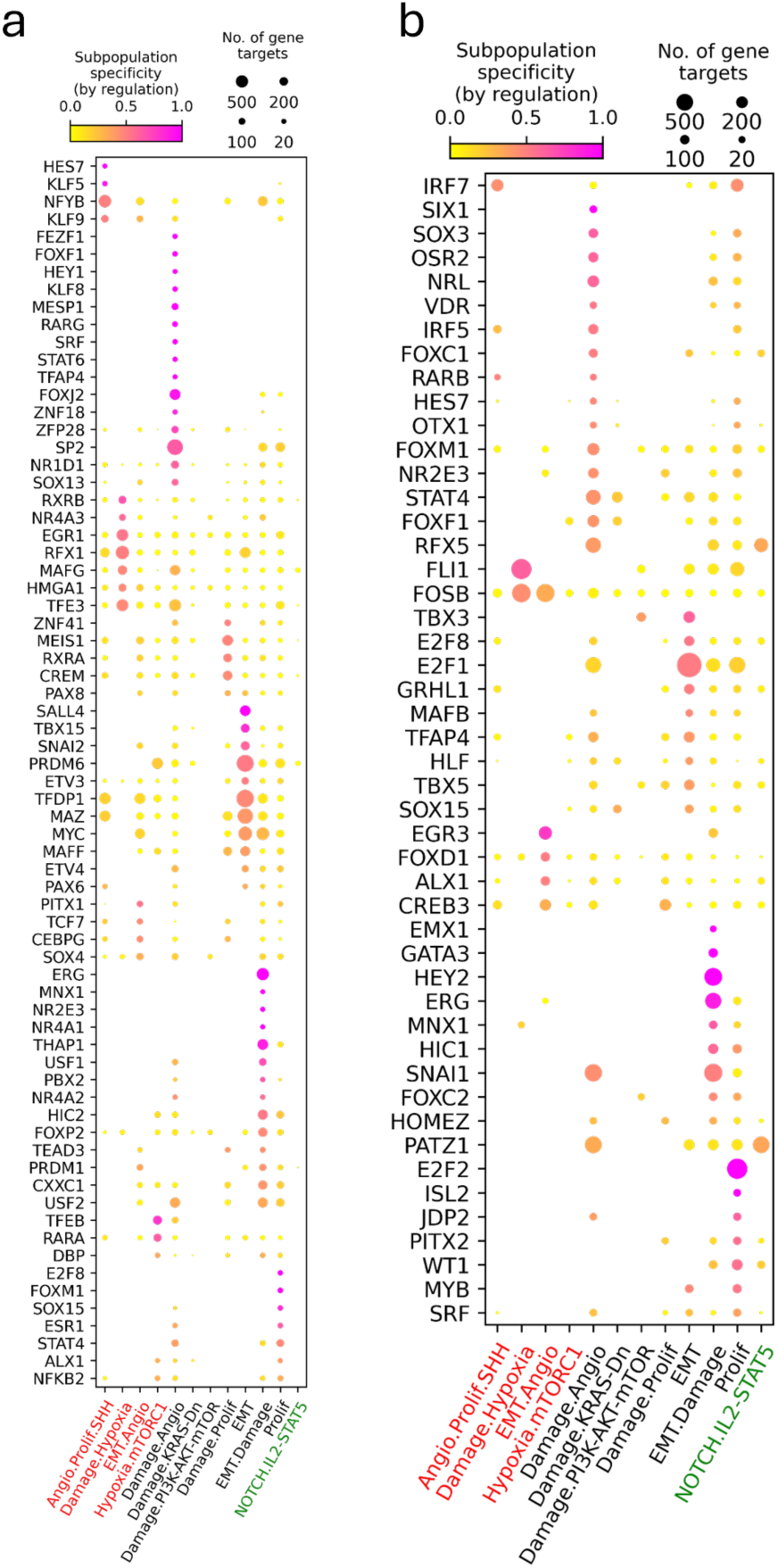
**a.** *Dictys* regulation markers for hypoxia, without replicate sample included. **b.** Regulation markers for samples across both conditions integrated.

**Supplementary Table 1.**
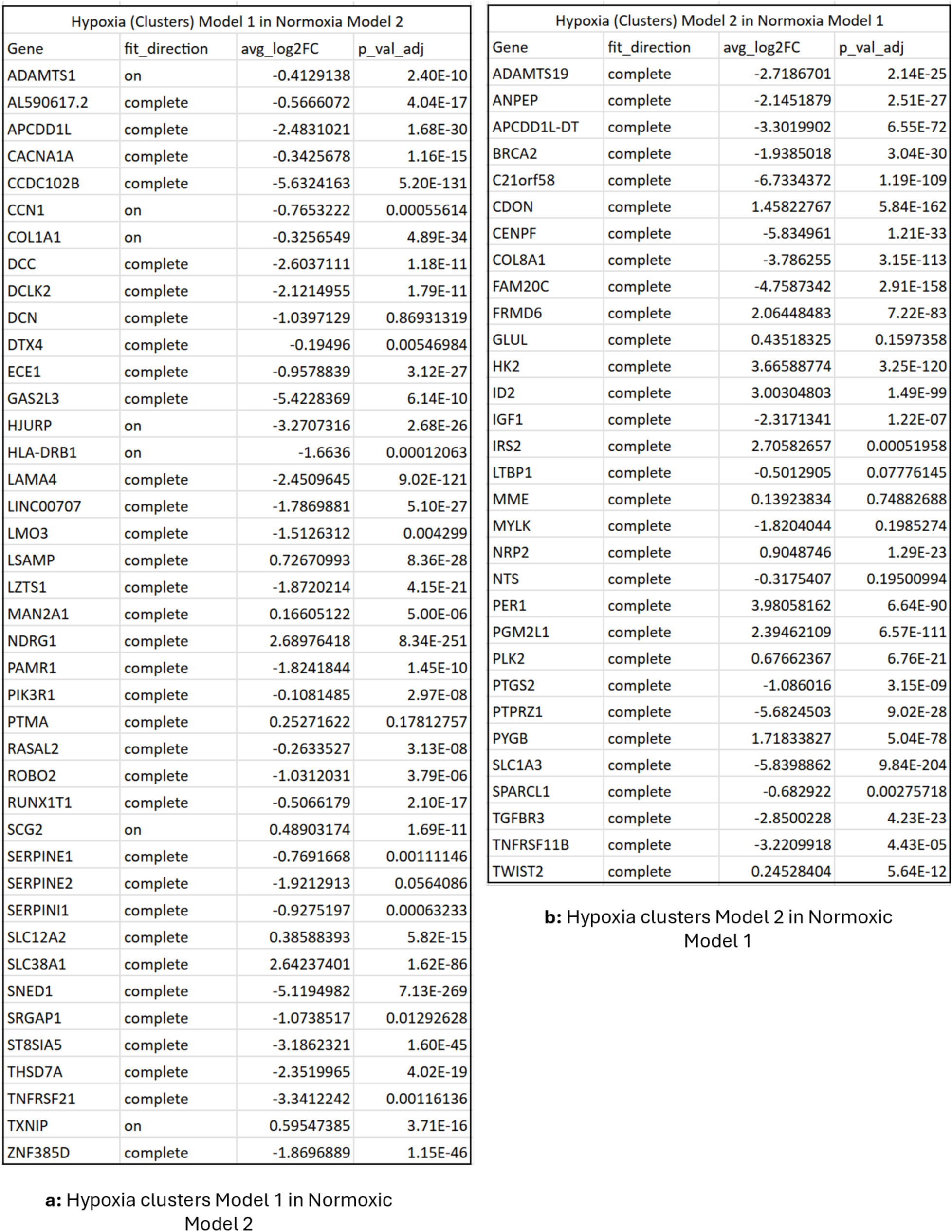
Models of Hypoxic clusters compared to hypoxic clusters in Normoxic sample.

**Supplementary Table 2.**
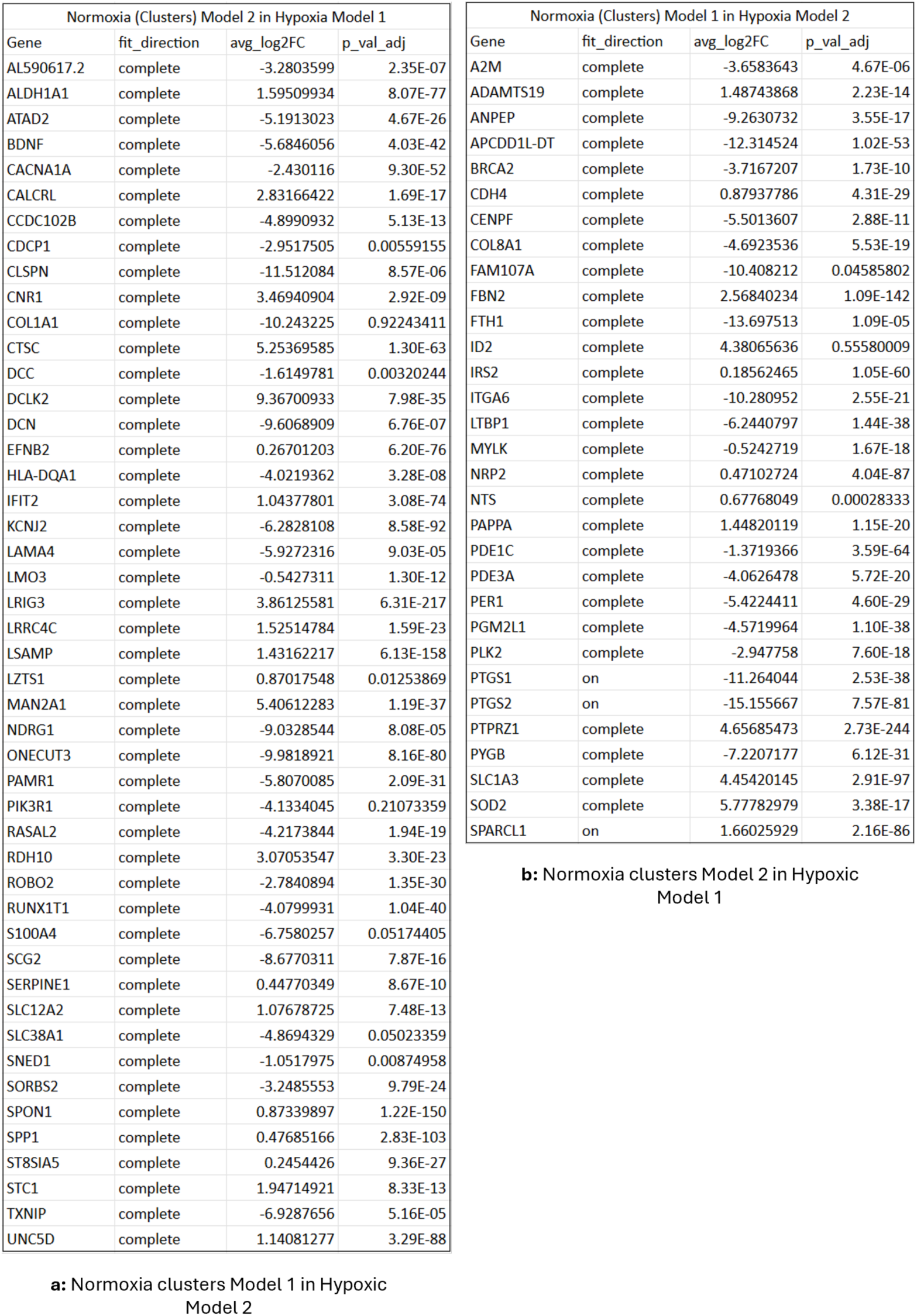
Models of Normoxic clusters compared to normoxic clusters in Hypoxic samples.

**Supplementary Table 3.**
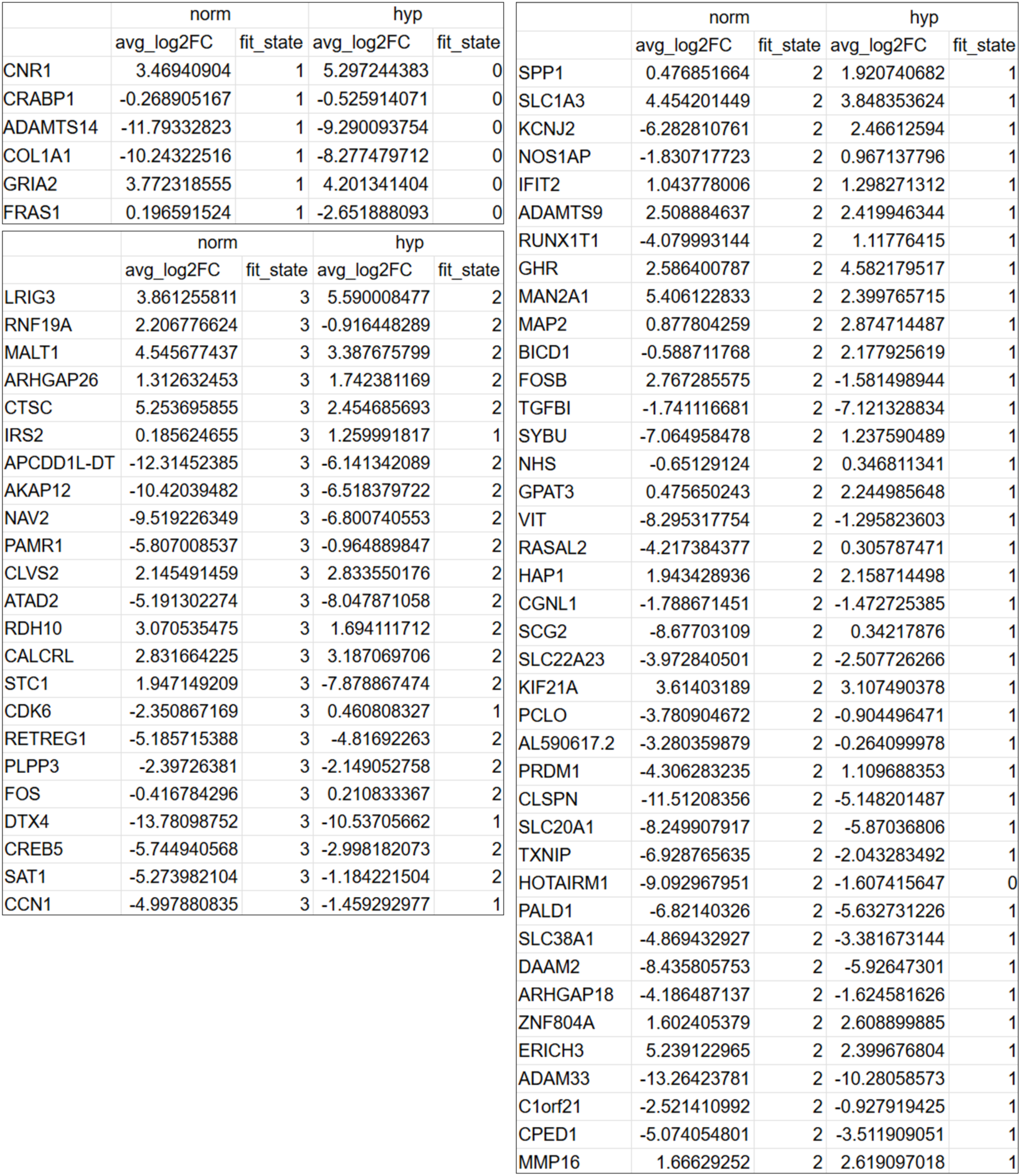
Normoxic fit states going in opposite direction of hypoxic sample.

**Supplementary Table 4.**
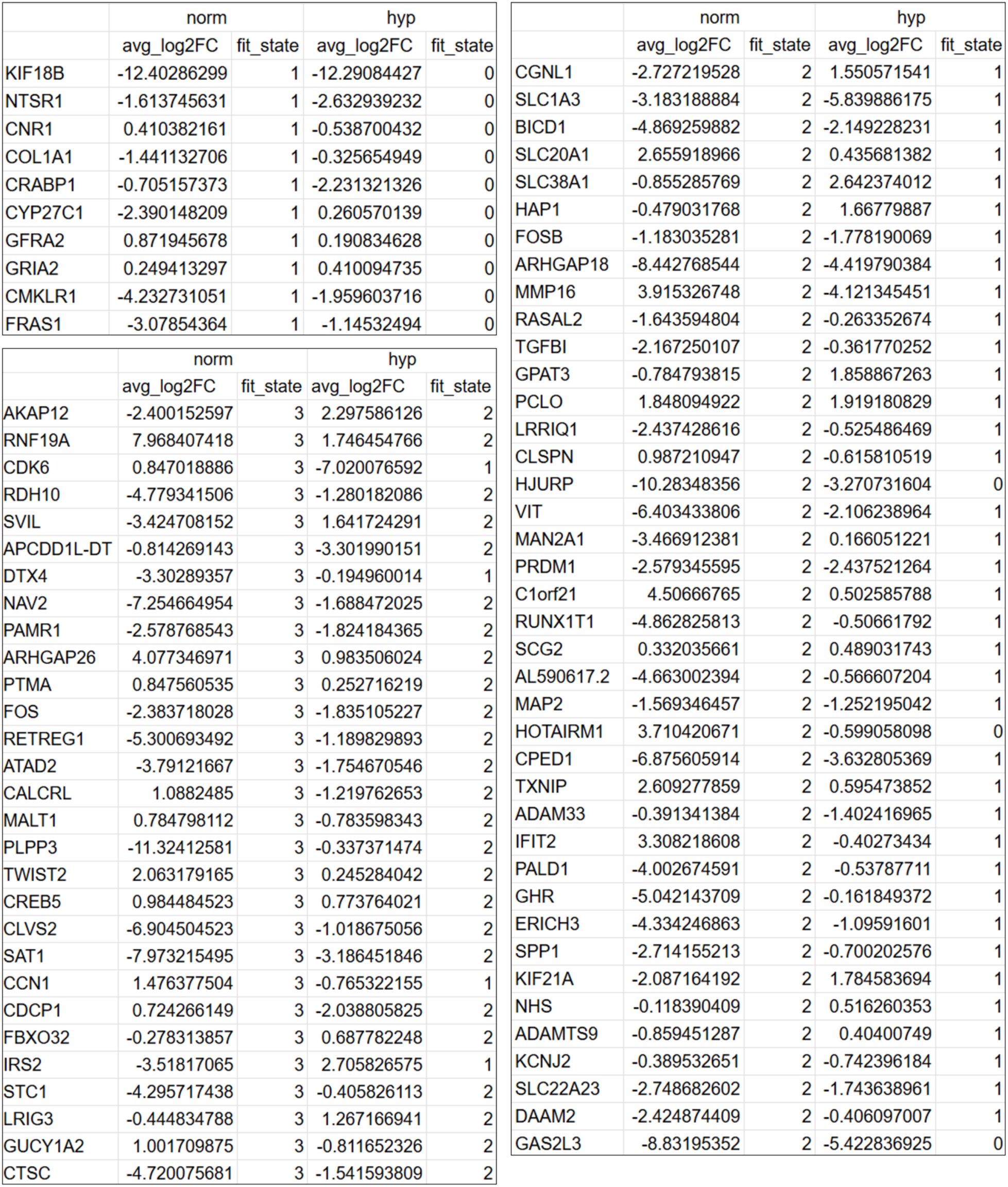
Hypoxic fit states going in opposite direction of normoxic sample.

## Methods

### Sample Preparation

#### Isolation and culture of primary GSCs

Collection of patient-derived glioblastoma tissue was approved by the IRB of Rhode Island Hospital. All patients provided written informed consent, with collections de-identified. Neurospheres were cultures from the tissue as previously described.^53^ GSCs used in this study were authenticated by ATCC using short tandem repeat analysis. The GSCs used were passage 18, cultures as attached on fibronectin-coated plates (10 g/mL), in a complete media containing 1X Neurobasal-A medium, B27-A supplement, 1 mg/mL Heparin, 20 ng/mL EGF and 20 ng/mL FGF.

#### Hypoxia

Cells were cultured on the same plate until last passage before nuclei isolation. These were then split onto two separate plates, and cultured for 3 days. Both plates were fed fresh media. The normoxic plate was placed back into the incubator. The hypoxic plate was placed into the hypoxia chamber (STEMCELL Technologies), with a plate of sterile water at the bottom for humidity, and oxygen was purged for 4 minutes. The hypoxia chamber was placed in the incubator, and the cells were cultured for 48 hours.

#### Nuclei Isolation

Nuclei were isolated with the buffers as listed the 10X Genomics Epi Multiome ATAC + Gene Expression kit protocol, with the addition of Lysis Dilution Buffer (10 mM Tris-HCL, pH 7.4; 10 mM NaCl; 3mM MgCl2; 1% BSA; nuclease-free water), and 1.5% BSA in Wash Buffer. All spins were with swinging bucket rotors. Cells were lifted from the plate using Accutase (STEMCELL Technologies) and spun with 5 mL Neurobasal-A at 230 xg for 5 mins. All media was aspirated, the cell pellet resuspended in 1 mL PBS + 2% BSA and transfered to 2 mL low adhesion microcentrifuge tubes. Cells were spun at 300 xg for 5 minutes. All supernatant was aspirated, 100 L of 0.3X lysis buffer was added, pipette mixed 10 times, and incubated on ice for 3 minutes. Then, 1 mL of chilled Wash Buffer was added, gently pipette mixed 5 times, and centrifuged at 500 xg for 5 minutes. Supernatant was aspirated, and wash step was repeated another 2 times. Nuclei were re-suspended in 100 µL PBS, counted using the automated Cell Countess, and then resuspended in according amounts of Diluted Nuclei Buffer.

#### Sequencing libraries

snATAC-seq and snRNA-seq libraries were generated using the Chromium Next GEM Single Cell Multiome Reagent Kit A, Chromium Next GEM Single Cell Multiome ATAC Kit A, Library Construction Kit B, Chromium Next GEM Chip J Single Cell Kit, Single Index Kit N Set A, and Dual Index Kit TT Set A, as per the Epi Multiome ATAC + Gene Expression protocol. The sequencing was done by the Molecular Biology Core Facilities at Dana-Fraber Cancer Institute using Illumina NovaSeq SP Flowcells (800M reads) for both ATAC and RNA, as well as the raw data processing, with CellRanger ARC counts outputs being provided.

### Analysis of snRNA-seq and sn-ATAC-seq

#### Multiome integration and cell exclusion

Following sequencing, we ended up with 8500/7500 cells, and 6500/6500 cells passed the sequencing threshold quality control for the hypoxia and normoxic condition samples, respectively. Once sequenced, one of the normoxic duplicates was found to have aberrant expression, and was excluded. Our three samples were read loaded as Seurat objects, using the filtered feature matrix from the gene expression data, and the ATAC peaks were added to the object using Signac. We then filtered out low quality cells, only including cells with: RNA counts between 3,000 and 90,000, ATAC counts between 3,000 and 60,000, TSS enrichment above 1, nucleosome signal below 2, and a mitochondrial genomic content below 20%. Final population was x/y, and z, for the two hypoxic samples and one normoxic sample, respectively.

#### Data anchoring and cluster determination

Samples had sn-ATAC-seq and sn-RNA-seq integrated in the previous step. Data was normalized using SCTransform, then performed latent semantic indexing to process DNA accessibility. Each of the samples were then anchored across each other, first the two hypoxic samples, then those to the normoxic sample. From here, the optimal clusters were determined via *scclusteval*, using the Jaccard similarity index,^14^ to be 100 for k.nn (equivalent of k.param for multiome data), 17 for the PCs, and a resolution of 1.7.

This provided us with 17 final clusters as seen in Fig. 1. To label each cluster according to functional pathway enrichment, the most highly expressed genes were first identified using FindMarkers. Then, the EnrichR platform allowed us to perform functional pathway enrichment analysis using the MSigDB Hallmark geneset, and labeled accordingly.

#### Transcription factor motif identification

Transcription factor-associated accessibility was identified using *chromVAR*^54^ and *motifmatchr*. Motifs were identified using Seurat’s FindMarkers for each cluster, with the logistic regression framework test (“LR”) used. The intersection of differentially expressed genes and differentially accessible motif peaks for each cluster was extracted, and scaled accordingly using min-max normalization set to [-1:1]. These were represented using *ComplexHeatmaps*.^55^ Transcription factor footprinting was also done

### Multiomic modeling

#### Building transcription factor networks

Transcription Factor regulatory networks were built using *Dictys*.^26^ Samples were ran in three different ways: independently, integrated across all three, and a final time with integrated hypoxic samples. The gene expression matrix was built from the data that already underwent cell exclusion (see above). We added unique suffixes to all BAM files and cell ID in the expression barcode matrix for the integrated samples, to account for any non-unique barcodes. The network inference was run on 2 GPUs with 12 cores. As quality control was already previously done, the networks were analyzed as described in the tutorial. We focused on TFs that were unique to glioblastoma for further analysis.

#### Multiome velocity and trajectory inference

Multiome velocity was analysed using velocyto, then MultiVelo. First, the bams were run through velocyto.R to generate the loom files the make up the spliced and unspliced data (with the “gex” removed from the file name). They were then analysed using MultiVelo. The two hypoxia samples were integrated, with all samples being filtered according to previous exclusion criteria. The data was then pre-processed, by selecting the top 5000 variable genes and filtering for minimum shared counts of at least 10, then the xy coordinates of the UMAP generated by MultiVelo was replaced with the xy coordinates of the Seurat UMAP with optimized clustering.

For further analysis, we compared the different outputs provided by MultiVelo. First, we extracted model 1 and model 2 genes from both the unique clusters and the entire sample, across both conditions, and searched for any intersection of opposite models for the same gene across samples. Similarly, we were able to extract all the genes that showed opposite behavior in the fit states between the two conditions over time, to identify any genes perturbed by the hypoxic microenvironment.

### RIDDLER

#### CNV Calling from sn-ATAC-seq

Single cell CNVs were called from fragment files using RIDDLER. RIDDLER was ran at 1MB window resolution, using default parameters and a total read filter of 10,000 reads per cell. No notable difference in output was observed from running RIDDLER on each batch separately vs all cells combined. CNV outputs from RIDDLER were then clustered using the Rphenograph package, using values of 20, 30, and 40 for parameter k and choosing the result with the highest modularity score. This resulted in 13 clusters, which we call the 13 CNV based subclones. Of these 13 clusters, clusters 10 through 13 consisted of less than 50 cells each, and were thus omitted from further subclone analysis.

#### Clone Enrichment

To test the relative enrichment or depletion of hypoxic and normoxic cells in each subclone, a chi-squared test was used. The table for the test consisted of the number of cells in the subclone for each condition in row 1, and the total number of cells in each condition in row 2. P-values for the tests were corrected using FDR, with values less than 0.05 taken as significant.

#### Differential CNV Detection

Differential CNVs between hypoxic and normoxic conditions were found in each subclone using a Wilcoxon Rank-Sum test. The test was performed individually for each 1MB window, testing the difference between the distribution of copy-numbers in each condition. Multiple hypothesis test correction for window p-values was performed using FDR, with a cutoff of 0.05. An additional cutoff was added to prune windows that did not have at least a 10% difference in the proportion of CNV events (i.e. copy-numbers that were non-diploid) between the two conditions.

## Data Availability

Code can be found here.

